# Botanical Medicines with Activity against Stationary Phase *Bartonella henselae*

**DOI:** 10.1101/2020.08.19.256768

**Authors:** Xiao Ma, Jacob Leone, Sunjya Schweig, Ying Zhang

**Author notes:** **Corresponding author**: Ying Zhang, MD, PhD.

## Abstract

*Bartonella henselae* is a Gram-negative, facultative intracellular bacterium which is the causative agent of cat scratch disease. In humans, infections with *B. henselae* can result in acute or chronic systemic infections with various clinical symptoms including local skin lesions, malaise, aches, chills, lymphadenopathy, endocarditis, or meningoencephalitis. The current treatment for *Bartonella* infections with antibiotics such as doxycycline and rifampin is not always effective presumably due to bacterial persistence. There have been various anecdotal reports of herbal extracts used for treating patients with persistent *Bartonella* infections but their activity on *B. henselae* is unknown. To test the potential antimicrobial activity of botanical or herbal medicines and develop better therapies for persistent *Bartonella* infections, in this study, we screened an herbal product collection against stationary phase *B. henselae in vitro* using SYBR Green I/ propidium iodide (PI) viability assay. These herbal medicines were selected by the fact that they are commonly used to treat Lyme and co-infections by patients and herbalists, and as a follow-up to our recent study where these herbs were tested against *B. burgdorferi*. We identified five herbal product extracts that had high activity against stationary phase *B. henselae* at 0.5% (*v/v*), including *Cryptolepis sanguinolenta*, *Juglans nigra*, *Polygonum cuspidatum*, *Scutellaria baicalensis*, and *Scutellaria barbata*. Among them, *Cryptolepis sanguinolenta*, *Juglans nigra*, and *Polygonum cuspidatum* could eradicate all stationary phase *B. henselae* cells within 7 days at 0.25% (*v/v*) in drug exposure time-kill assay, whereas *Scutellaria baicalensis* and *Scutellaria barbata* showed relatively poor activity. The minimum inhibitory concentration (MIC) determination of these top hits indicated they were not only active against stationary phase non-growing *B. henselae* but also had good activity against log phase growing *B. henselae*. Our findings may help to develop more effective treatments for persistent *Bartonella* infections.

## INTRODUCTION

*Bartonella* species are fastidious Gram-negative facultative intracellular pathogens with a unique intraerythrocytic lifestyle, which usually require an obligately bloodsucking arthropod vector and a mammalian host during their life cycle [1]. At least thirteen *Bartonella* species are known to be pathogenic in humans, leading to either acute or chronic infections with known diseases such as cat scratch disease, trench fever, Carrion’s disease, and bacillary angiomatosis [2]. Infection typically begins with cutaneous inoculation and the pathogenesis of *Bartonella* involves establishing a primary niche in the endothelium [3] before seeding into the blood stream leading to intraerythrocytic infection [4–6]. *Bartonella* has been described as a “stealth pathogen” [7] due to its ability to cause persistent bacteremia, evade the immune system, and cause varying types and severity of symptomatology [6–10]. *Bartonella* can induce vasoproliferative tumor formation [11] and a recent review of the public health implications of bartonellosis discussed the need to research *Bartonella*’s potential role in breast cancer tumorigenesis [12]. Preliminary data have also implicated bartonellosis in neuropsychiatric manifestations including Autism Spectrum Disorder [13], homicidality [14], and hallucinations [15].

*Bartonella henselae* is the most common causative agent of cat scratch disease, with symptoms of malaise, decreased appetite, aches, headache, chills, arthritis, and lymphadenopathy that can persist for several months [16]. In certain situations, disseminated *B. henselae* may lead to the development of serious complications including ocular manifestations including neuroretinitis [17], neurologic manifestations including encephalopathy, seizures, cerebral vasculitis, meningoencephalitis [18][19][20], visceral organ involvement including hepatomegaly and/or splenomegaly [21], and cardiac manifestations including endocarditis [2][22], which can have a particularly high mortality [18]. Case studies have also implicated *B. henselae* in monoclonal and biclonal gammopathy [23] and various auto-immune manifestations including pediatric acute-onset neuropsychiatric syndrome (PANS) [24], transverse myelitis [25,26], and autoimmune thyroiditis [27].

Emerging evidence suggests *B. henselae* may also serve as a co-infective pathogen in patients with other vector-borne diseases [12]. In particular, ticks are known to be polymicrobially infected with multiple pathogens [28] including *Borrelia burgdorferi,* the most common etiologic agent in Lyme disease. Although the ability of ticks to serve as a competent vector for transmitting *B. henselae* to humans is debated, ticks have been documented to carry *B. henselae* [12] and case reports link *B. henselae*-related disease following a tick attachment [29–31]. Multiple papers document *B. henselae* exposure in patients with Lyme disease and patients with co-infections may experience more severe and protracted clinical manifestations [31–34].

Currently there is no single treatment effective for all *Bartonella*-associated diseases, with different antibiotics recommended for use according to different presentations [35]. The first-line antibiotics for the treatment of *Bartonella*-associated infection include doxycycline, erythromycin, azithromycin, gentamicin, rifampin, ciprofloxacin, and tetracycline as well as some drug combinations such as doxycycline plus gentamicin or doxycycline plus rifampin [35][36]. However, the recommended treatment of systemic *B. henselae* infections is based on limited trial data [37] with various treatment failures, probably related to antibiotic resistance and bacterial persistence [36][38]. Therefore, identifying novel antimicrobials targeting persistent *Bartonella* pathogens could significantly assist in development of improved therapeutic protocols for the treatment of *Bartonella*-associated diseases.

Botanical medicines have a long history of documented use since ancient Mesopotamia, ancient China and ancient India. The safety and efficacy of these medicines is supported by centuries of use in various traditional medicine systems such as Ayurveda and Traditional Chinese Medicine [39–41]. Several recent reports have concluded that the frequency of severe adverse effects related to the use of botanical medicine is rare [42–44]. Unlike conventional antibiotics, which can have a detrimental impact on the microbiome and increase microbial resistance [45, 46], botanical medicines may have a beneficial impact on the microbiome [47]. Many botanical extracts have been reported to have antimicrobial activities, for example, *Laserpitium zernyi* herb extracts were active against different bacterial species including *Pseudomonas aeruginosa*, *Micrococcus luteus*, *Enterococcus faecalis* and *Bacillus subtilis*, and different extracts of *Ononis arvensis* showed antimicrobial activity against *Escherichia coli*, *P. aeruginosa*, *Salmonella typhimurium*, *Staphylococcus aureus* and *Candida albicans*. Some Mediterranean herb extracts could inhibit growth of representative oral microorganisms and biofilm formation of *Streptococcus mutans* [48–50]. Members of our group previously documented 32 essential oils derived from botanical medicines that were found to have better activity against stationary phase *B. henselae* compared to antibiotic controls [51]. It is becoming increasingly important to study botanical medicines with potential antimicrobial activity for improved treatment of infections due to antibiotic resistant bacteria. According to the World Health Organization, antibiotic resistance is currently “one of the biggest threats to global health, food security, and development” [52].

Previously, we have developed a rapid high-throughput drug screen method with the help of SYBR Green I/ propidium iodide (PI) viability assay, leading to successful identification of various active drugs as well as drug combinations against stationary phase *Borrelia burgdorferi* as a surrogate model of persister bacteria [53–56]. In two recent studies, we screened the FDA-approved drug library and a collection of essential oils using this method to identify active drug candidates with potential for treating *Bartonella* infections [51][57]. In this current study, we adapted the same methodology to screen an herbal product collection which we used recently on *Borrelia burgdorferi* [56] and successfully identified various botanical extracts with activity against stationary phase *B. henselae*, which has successfully been used as a model of persister drug screens [53–56] because of its higher content of persister cells than the log phase culture [58]. The implications of these findings for better therapy of persistent *Bartonella*-associated infections are discussed.

## MATERIALS AND METHODS

### Bacterial strain, culture media and culture conditions

*Bartonella henselae* JK53 strain was obtained from BEI Resources (ATCC), NIAID, NIH. *B. henselae* JK53 was cultured in Schneider’s Drosophila medium (Life Technologies Limited, Paisley, UK) supplemented with 10% fetal bovine serum (FBS) (Sigma-Aldrich, Co., St. Louis, MO, USA) and 5% sucrose (Fisher Scientific, New Jersey, USA) as described previously [57][59]. Cultures were incubated in sterile 15 mL or 50 mL polypropylene conical tubes (Corning, New York, USA) in microaerophilic incubator without shaking at 37 °C, 5% CO_2_. Based on our previous study, the one-day-old and five-day-old cultures were considered as log phase and stationary phase, respectively [57]. The Columbia sheep blood agar (HemoStat Laboratories, Dixon, CA, USA) was used to perform the colony count in drug exposure assay, which was also cultured at 37 °C, 5% CO_2_.

### Drugs and herbal products

A panel of herbal products was purchased from KW Botanicals Inc. (San Anselmo, CA, USA) and Heron Botanicals (Kingston, WA, USA) [56]. These botanical products were chosen based on significant antimicrobial activity against other bacterial pathogens shown by previous studies [60–64], anecdotal clinical usage reported by patients, favorable safety profiles, and the ability to be absorbed systemically. Most of the botanicals were identified via macroscopic and organoleptic methods with voucher specimens on file with the respective production facilities. Additional details on sourcing, testing and validation of botanical and natural medicines used are summarized in Table S1. Most botanical extracts were provided as alcohol extracts in 30%, 60%, and 90% alcohol dilutions, and the alcohol used was also tested separately as a control in the same dilutions of 30%, 60%, and 90%. Herbal products were dissolved in dimethyl sulfoxide (DMSO) at 5% (*v/v*), followed by dilution at 1:5 into five-day-old stationary phase cultures to achieve 1% final concentration. To make further dilutions for evaluating anti-*Bartonella* activity, the 1% herbal products were further diluted with the same stationary phase cultures to achieve desired concentrations. Antibiotics including azithromycin (AZI), daptomycin (DAP), doxycycline (DOX), gentamicin (GEN), methylene blue, miconazole, and rifampin (RIF) included as controls were purchased from Sigma-Aldrich (USA) and dissolved in appropriate solvents [65] to form 10 mg/mL stock solutions. All the antibiotic stocks were filter-sterilized by 0.2 μm filters except the DMSO stocks and then diluted and stored at −20 °C.

### Microscopy techniques

*B. henselae* cultures treated with different herbal products or control drugs were stained with SYBR Green I (10 × stock, Invitrogen) and propidium iodide (PI, 60 μM, Sigma) mixture dye and then examined with BZ-X710 All-in-One fluorescence microscope (KEYENCE, Inc., Osaka, Japan). The SYBR Green I/PI pre-mixed dye was added to the sample in a 1:10 ratio of the dye against the sample volume and mixed thoroughly, followed by incubating in the dark at room temperature for 15 minutes. SYBR Green I is a green permeant dye that stains all cells whereas propidium iodide (PI) as an orange-red impermeant dye stains only dead or damaged cells with a compromised cell membrane. Thus, live cells with intact membrane will be stained only by SYBR Green I as green cells, while damaged or dead cells with a compromised cell membrane will be stained orange-red by PI. The residual bacterial viability could then be assessed by calculating the ratio of green/red fluorescence, respectively, as described previously [53]. The stained samples were confirmed by analyzing three representative images of the same bacterial cell suspension using fluorescence microscopy. BZ-X Analyzer and Image Pro-Plus software were used to quantitatively determine the fluorescence intensity.

### Screening of herbal product collection against stationary phase *B. henselae* using SYBR Green I/PI viability assay

For the primary herbal products screening, each product was assayed in two concentrations, 1% (*v/v*) and 0.5% (*v/v*). A five-day-old stationary phase *B. henselae* culture was used for the primary screen. Firstly, 40 μL 5% herbal product DMSO stocks were added to 96-well plates containing 160 μL *B. henselae* culture, respectively, to obtain the desired concentration of 1%. Then the 0.5% concentration was obtained by mixing 100 μL 1% treatment with 100 μL *B. henselae* culture. Antibiotics including AZI, DAP, DOX, GEN, methylene blue, miconazole and RIF were used as control drugs at their Cmax. Cmax is a pharmacokinetic measure referring to the maximum serum concentration that a drug achieves in a specified compartment such as blood after the drug has been administered. Control solvents including DMSO, 30%, 60%, and 90% alcohol were also included. Plates were sealed and placed in a 37°C incubator without shaking over a period of three days. SYBR Green I/ PI viability assay was used to assess the live/dead cell ratios after drug exposure as described [53]. Briefly, the SYBR Green I/PI dye was added to the sample followed by incubation in the dark for 15 minutes. Then the plate was read by microplate reader (HTS 7000 plus Bioassay Reader, PerkinElmer Inc., USA). The green/red (538 nm/650 nm) fluorescence ratio of each well was used for calculating the residual viability percentage, according to the regression equation of the relationship between residual viability percentage and green/red fluorescence ratio obtained by least-square fitting analysis as described [57]. All tests were run in triplicate.

### Drug exposure assay of active hits

The active hits from the primary screens were further confirmed by drug exposure assay. The five-day-old stationary phase *B. henselae* culture was used for drug exposure experiments and was treated with 0.25% (*v/v*) active herbal products respectively. Control antibiotics including AZI, DAP, DOX, GEN, methylene blue, miconazole and RIF were used at their Cmax. Control solvents were also included. The drug exposure assay was carried out in 15 mL polypropylene conical tubes over the course of seven days at 37°C, 5% CO_2_ without shaking. At each time point, a portion of bacterial cells was collected by centrifugation and rinsed twice with fresh Schneider’s medium followed by re-suspension in fresh Schneider’s medium. Then the cell suspension was serially diluted and plated on Columbia blood agar plates for viable bacterial counts (colony forming unit, CFU). The plates were incubated at 37 °C, 5% CO_2_ until visible colonies appeared and the CFU/mL was calculated accordingly. All tests were run in triplicate.

### Minimum inhibitory concentration (MIC) determination of active hits

The standard microdilution method was used to measure the minimum inhibitory concentration (MIC) of each herbal product needed to inhibit the visible growth of *B. henselae* after a five-day incubation period as described [51][57]. The diluted one-day-old *B. henselae* log phase culture was used for MIC determination. The 5% herbal product stocks were added into 96-well plates containing 1×10^6^ bacteria per well with fresh modified Schneider’s medium, respectively, to achieve 1% final concentration. Other lower concentrations were obtained by mixing 1% treatment with diluted one-day-old log phase *B. henselae* culture. Plates were sealed and incubated at 37 °C without shaking for five days. Then the bacterial cell proliferation was assessed using the SYBR Green I/PI assay and the bacterial counting chamber. The MIC is the lowest concentration of the herbal product that prevented the visible growth of *B. henselae*. All tests were run in triplicate.

### Statistical analysis

The statistical analysis was performed using two-tailed Student’s *t*-test. Mean differences were considered statistically significant if P value was < 0.05. Analyses were performed using Image Pro-Plus, GraphPad Prism, and Microsoft Office Excel.

## RESULTS

### Screening the herbal product collection to identify herbs active against non-growing stationary phase *B. henselae*

In this study, we adapted the SYBR Green I/PI viability assay developed previously [51][57] to identify herbal products with significant activity against stationary phase *B. henselae* compared to antibiotic controls. As described above, we tested a panel of botanical extracts and their corresponding controls against a five-day-old stationary phase *B. henselae* culture in 96-well plates incubated for 3 days. For primary screens, all the herbal products were applied at two concentrations, 1% (*v/v*) and 0.5% (*v/v*), respectively. Meanwhile, the currently used antibiotics for bartonellosis treatment such as doxycycline, azithromycin, gentamicin, and rifampin were included as control drugs for comparison, as well as the previously identified FDA-approved drugs effective against *B. henselae* including daptomycin, methylene blue and miconazole [57] (Table 1). All of the pharmaceutical antibiotics were used at their Cmax. In the primary screens, four different alcohol extracts of *Juglans nigra*, three different alcohol extracts from *Cryptolepis sanguinolenta*, one alcohol extract of *Polygonum cuspidatum*, one glycerite extract of *Polygonum cuspidatum*, one glycerite extract of *Scutellaria baicalensis* (huang qin), and one glycerite extract of *Scutellaria barbata* (ban zhi lian) at both 1% (*v/v*) and 0.5% (*v/v*) were found to have strong activity against stationary phase *B. henselae* compared to the control antibiotics including AZI, DOX, GEN, and RIF according to the plate reader results (Table 1). In contrast, *Andrographis paniculate, Stevia rebaudiana, Artemisia annua, Uncaria rhynchophylla, Uncaria tomentosa, Rhizoma coptidis, Citrus paradise, Dipsacus fullonum, Campsiandra angustifolia, Otoba parvifolia* and colloidal silver did not show significant activity against stationary phase *B. henselae* (Table S2). Therefore, the following active herb extracts were selected as top hits, including *Juglans nigra* 30%, 45%, 60%, and 90% alcohol extract, *Polygonum cuspidatum 30%* alcohol extract, *Polygonum cuspidatum* glycerite extract, *Cryptolepis sanguinolenta* 30%, 60%, and 90% alcohol extract, *Scutellaria baicalensis* glycerite extract, and *Scutellaria barbata* glycerite extract. These top hits were chosen based on the fact that a lower percentage of viable cells remained compared to the other natural antimicrobials used in this study as well as exhibiting significantly better activity compared to the current antibiotics used to treat *Bartonella* infections, including AZI, DOX, GEN, and RIF, as indicated by the statistically analysis that the P value was < 0.05. Our previous experience showed some compounds in the herbal extracts could interfere with the SYBR Green I/PI assay due to their autofluorescence. To eliminate this impact, we checked the residual cell viability by examining microscope images of the herbal extract treated samples to confirm the plate reader results. As shown by fluorescence microscopy, solvents such as DMSO and alcohol did not have significant impact on residual bacterial cell viability compared to the drug free control (Figure 1 and Table 1), as the P value was > 0.05. Clinically used antibiotics against *Bartonella* infections such as AZI and DOX only showed weak activity when used at their Cmax (residual viability above 60%) (Table 1). Antibiotics reported to have a clinical improvement for *Bartonella* infections including GEN and RIF showed relatively better activity (residual viability below 50%) against stationary phase *B. henselae* than AZI and DOX. FDA-approved drugs that we identified as effective against stationary phase *B. henselae* (DAP, methylene blue, and miconazole) [57] had better activity (residual viability below 40%) than the other antibiotics tested. Among the five top herbal hits that had better activity (residual viability between 0% and 16%) against stationary phase *B. henselae* than most control antibiotics, the most active herbal products were *Juglans nigra* and *Polygonum cuspidatum* alcohol extracts of different concentrations. However, the fluorescence microscope observation of *Juglans nigra* 30% and 45% alcohol extracts, and *Polygonum cuspidatum* glycerite extract at 0.5% treatment exhibited significantly higher percentage of green (live) cells compared with the plate reader results (Figure 1 and Table 1), as indicated by the statistically analysis that the P value was < 0.05, which were also higher than that of most control antibiotics, indicating the relatively poor accuracy of the plate reader results and poor activity of these herbal products at these particular concentrations. Therefore, they were excluded from active hits for subsequent MIC testing and drug exposure assay (see below). Alcohol extracts from *Cryptolepis sanguinolenta* of different concentrations also exhibited strong activity against stationary phase *B. henselae* as shown by red (dead) cells in fluorescence microscope observation (Figure 1), which is consistent with the plate reader results. Glycerite extracts from the two *Scutellaria* plants, including *Scutellaria baicalensis* (huang qin) and *Scutellaria barbata* (ban zhi lian), also showed good activity with low percentages of residual viable bacterial cells remaining (Figure 1).

**Table 1.** Activity of top active herbal products against stationary phase *B. henselae* ^1^.

| Herbal Products (HP) and Control Drugs | Trade Names of Herbal Products | Residual Viability (%) after 1% HP or Antibiotic Treatment |  | Residual Viability (%) after 0.5% HP Treatment |  |
| --- | --- | --- | --- | --- | --- |
|  |  | Plate Reader<sup>2</sup> | Microscope<sup>3</sup> | Plate Reader<sup>2</sup> | Microscope<sup>3</sup> |
| Drug free control |  | 70% | 75% |  |  |
| DMSO control |  | 60% | 70% |  |  |
| 30% alcohol |  | 70% | 78% |  |  |
| 60% alcohol |  | 80% | 82% |  |  |
| 90% alcohol |  | 65% | 63% |  |  |
| AZI |  | 70% | 65% |  |  |
| DOX |  | 66% | 60% |  |  |
| GEN |  | 44% | 50% |  |  |
| RIF |  | 31% | 45% |  |  |
| DAP |  | 6% | 10% |  |  |
| Methylene Blue |  | 29% | 35% |  |  |
| Miconazole |  | 40% | 50% |  |  |
| *Juglans nigra* 45% AE<sup>5</sup> | Hu tao ren | 0%<sup>4</sup> | 0% | 9% | 59% |
| *Juglans nigra* 60% AE<sup>5</sup> | Black walnut | 0%<sup>4</sup> | 0% | 4% | 32% |
| *Juglans nigra* 90% AE<sup>5</sup> | Black walnut | 0%<sup>4</sup> | 7% | 6% | 30% |
| *Juglans nigra* 30% AE<sup>5</sup> | Black walnut | 2% | 2% | 6% | 63% |
| *Polygonum cuspidatum* 30% AE<sup>5</sup> | Japanese knotweed | 6% | 0% | 8% | 21% |
| *Polygonum cuspidatum* GE<sup>6</sup> | Hu zhang | 8% | 18% | 13% | 70% |
| *Cryptolepis sanguinolenta* 30% AE<sup>5</sup> | Cryptolepis | 8% | 10% | 14% | 42% |
| *Scutellaria baicalensis* GE<sup>6</sup> | Huang qin | 8% | 12% | 11% | 25% |
| *Scutellaria barbata* GE<sup>6</sup> | Ban zhi lian | 8% | 50% | 2% | 48% |
| *Cryptolepis sanguinolenta* 60% AE<sup>5</sup> | Cryptolepis | 10% | 24% | 14% | 34% |
| *Cryptolepis sanguinolenta* 90% AE<sup>5</sup> | Cryptolepis | 11% | 25% | 14% | 47% |

<sup>1</sup> A five-day-old stationary phase *B. henselae* culture was treated with herbal products (1% or 0.5%) (v/v) or control drugs for three days. Drug concentrations used in this experiment were based on their Cmax and were as follows: 0.2 µg/mL AZI, 2.4 µg/mL DOX, 10 µg/mL GEN, 7.8 µg/mL RIF, 60 µg/mL DAP, 2.9 µg/mL methylene blue, and 6.3 µg/mL miconazole. <sup>2</sup> Residual viability was calculated according to
the regression equation and the ratio of Green/Red fluorescence obtained by SYBR Green I/PI assay.<sup>3</sup> Residual viability was assayed by fluorescence microscope counting.<sup>4</sup> Values of SYBR Green I/PI calculated by the plate reader were lower than 100% dead cells.<sup>5</sup> Abbreviation: AE: alcohol extract; GE: glycerite extract.

**Figure 1.**
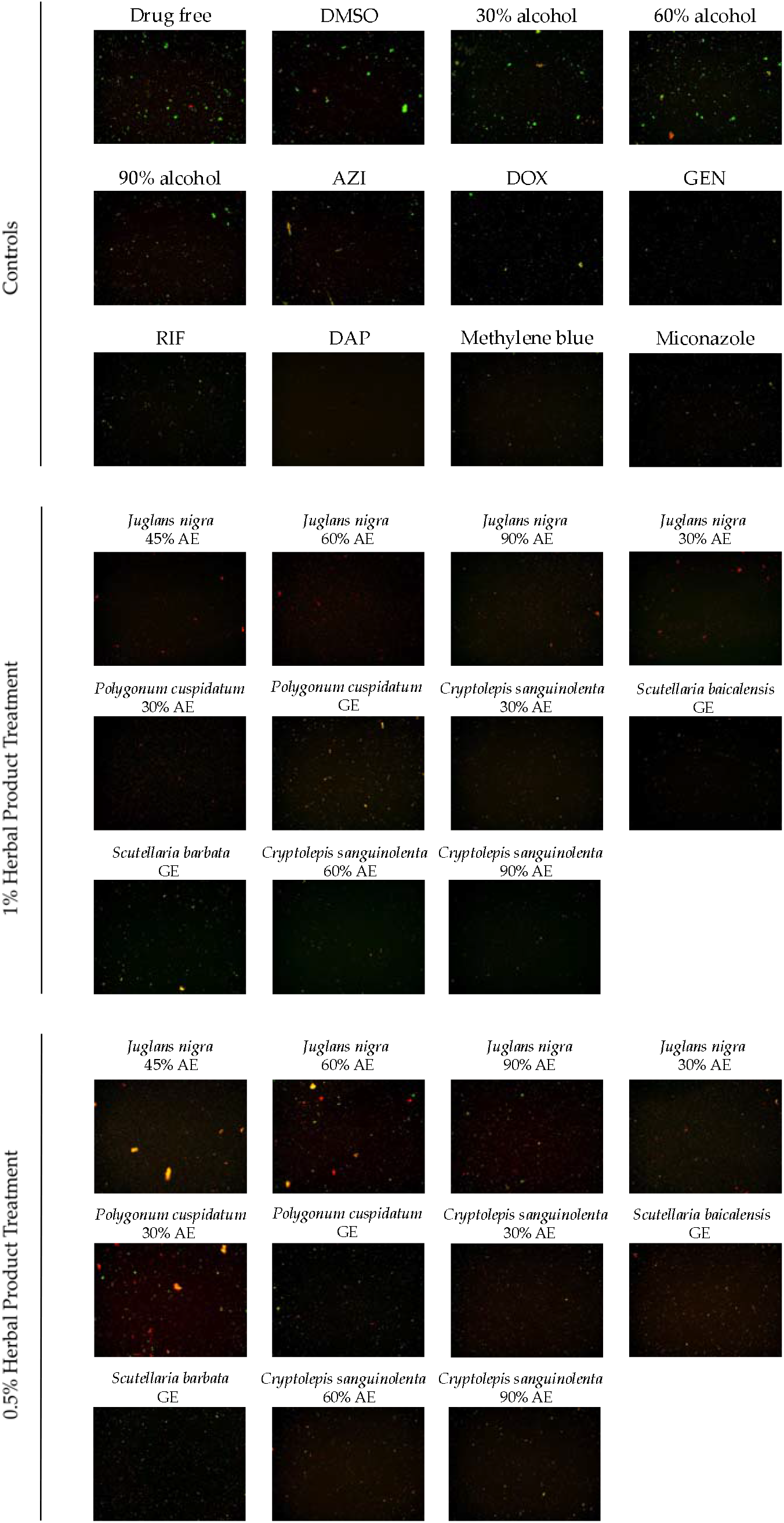
Effect of top hits of herbal products against stationary phase *B. henselae* in comparison with control drugs. A five-day-old stationary phase *B. henselae* culture was treated with 1% (*v/v*) or 0.5% (*v/v*) herbal products or control antibiotics for three days followed by SYBR Green I/PI viability assay and fluorescence microscopy (400 × magnification). Drug concentrations used were based on their Cmax and were as follows: 0.2 μg/mL AZI, 2.4 μg/mL DOX, 10 μg/mL GEN, 7.8 μg/mL RIF, 60 μg/mL DAP, 2.9 μg/mL methylene blue, and 6.3 μg/mL miconazole. Green cells represent live cells and red cells represent dead cells.

### Time-kill curves of active hits

To further demonstrate the efficacy of the active herbal products identified from the primary screens in eradicating persistent *B. henselae* cells, we performed a time-kill drug exposure assay against a five-day-old stationary phase *B. henselae* culture at a lower concentration of 0.25% (*v/v*), along with their corresponding solvent controls. Meanwhile, clinically used antibiotics to treat *Bartonella* infections including AZI, DOX, GEN, and RIF were used at their Cmax as controls. Compared to the drug free control, as shown in Figure 2 and Table 2, some clinically used antibiotics such as AZI and DOX showed poor activity in killing stationary phase *B. henselae* partly due to their low Cmax. Other antibiotics such as GEN and RIF exhibited better activity which could eradicate all *B. henselae* cells by day 7 and day 5, respectively. The difference of residual viabilities of stationary phase *B. henselae* after treatment by control solvents and without drug treatment was not statistically significant, as the P value was > 0.05. All three *Cryptolepis sanguinolenta* alcohol extracts of different concentrations were able to eradicate all *B. henselae* cells in the seven-day drug exposure, where *Cryptolepis sanguinolenta* 60% alcohol extract was the most active herbal product that killed *B. henselae* with no detectable CFU after five-day exposure. *Juglans nigra* in 60% and 90% alcohol extracts both exhibited good activity that eradicated all *B. henselae* cells without viable cells being recovered after the seven-day drug exposure. *Polygonum cuspidatum* 30% alcohol extract was also effective to kill all *B. henselae* cells by day 7. However, *Scutellaria barbata* (ban zhi lian) and *Scutellaria baicalensis* (huang qin) showed poor activity at the concentration of 0.25% (*v/v*) during this seven-day drug exposure, with considerable numbers of viable cells remaining after treatment.

**Figure 2.**
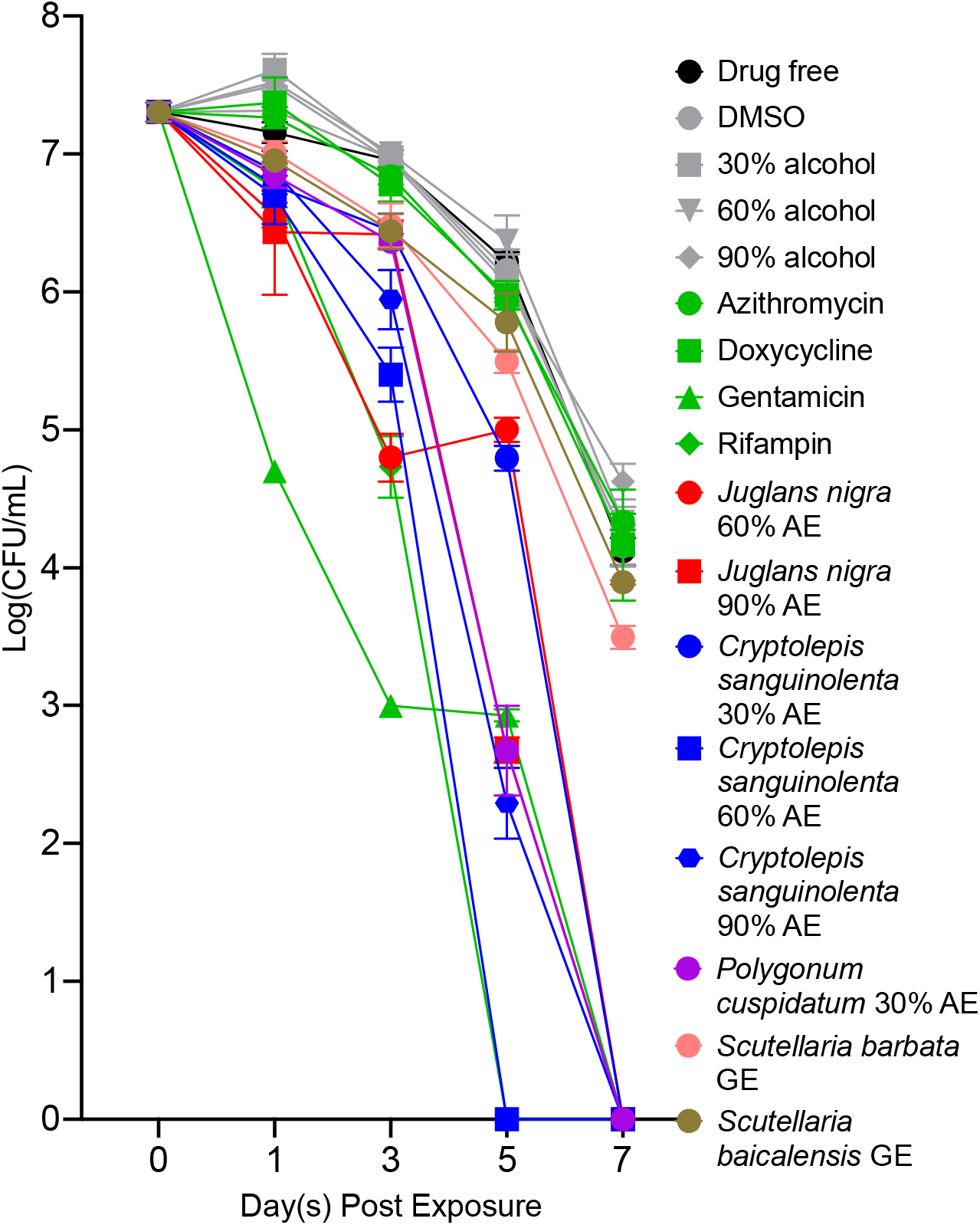
Time-kill curves of active herbal product treatment against stationary phase *B. henselae* in comparison with control drugs. The herbal products or control antibiotics were added to the five-day old stationary phase culture respectively at time point 0, and at different times of drug exposure (day 1, day 3, day 5, and day 7), portions of bacteria were removed and washed and plated on Columbia blood agar plates for CFU counts. The herbal product concentration used in this experiment was 0.25% (*v/v*). Drug concentrations used in this experiment were based on their Cmax and were as follows: 0.2 μg/mL AZI, 2.4 μg/mL DOX, 10 μg/mL GEN, and 7.8 μg/mL RIF.

**Table 2.**
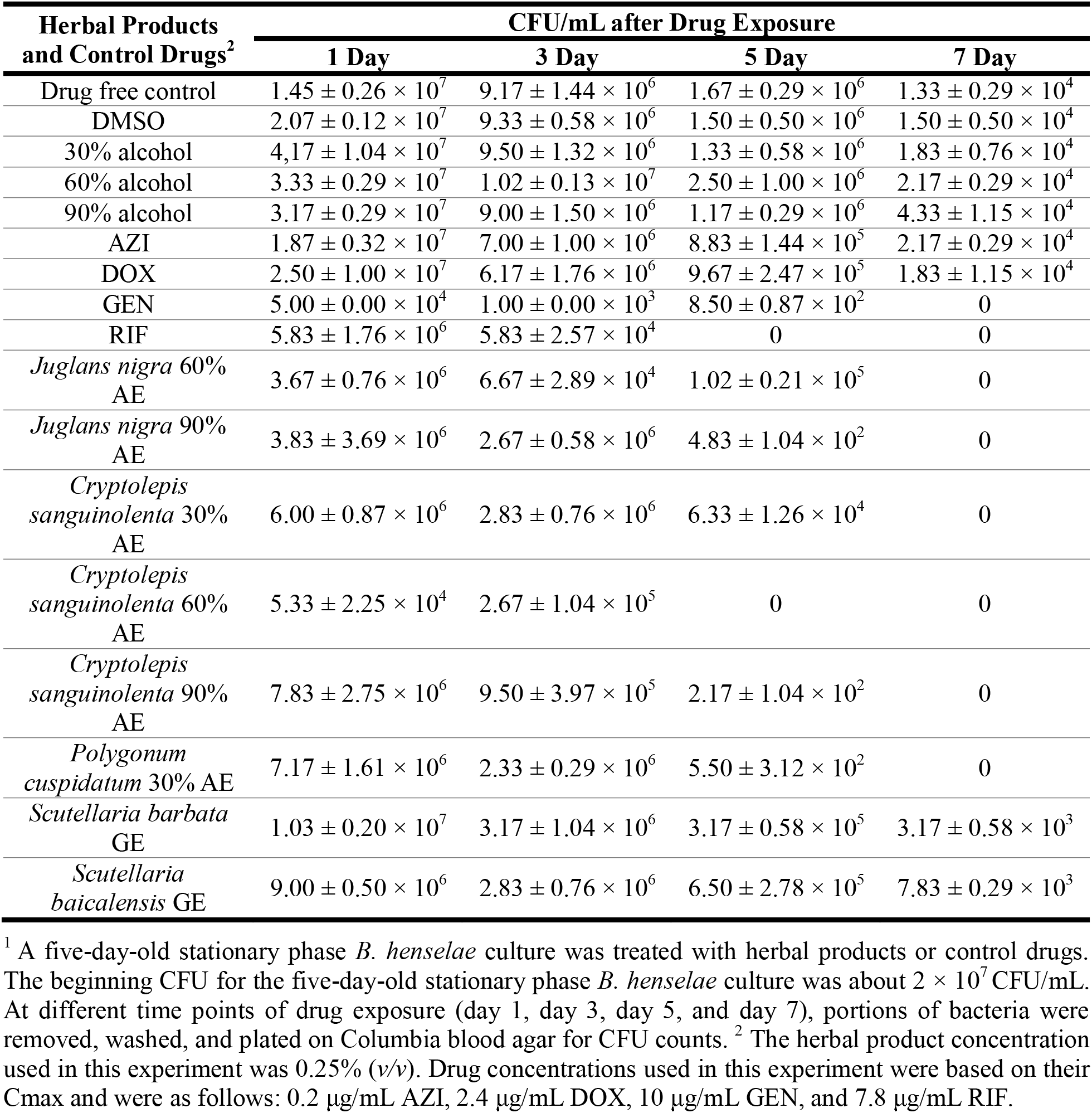
Drug exposure assay of top active herbal products against *B. henselae* stationary phase culture^1^.

### Minimum inhibitory concentration (MIC) determination of active hits

The activity of antibiotics against non-growing bacteria is not always correlated with that against growing bacteria [57]. Thus, it was also necessary to determine the MICs of these active herbal products against log phase growing *B. henselae*. The MIC determination of herbal products for *B. henselae* was conducted by the standard microdilution method as described [51][57]. As shown in Table 3, *Juglans nigra* 60% extract was the most active herbal product among the top 5 hits, capable of inhibiting visible *B. henselae* proliferation at 0.125%-0.25% (*v/v*). Other herbal products including *Juglans nigra* 90% alcohol extracts, *Polygonum cuspidatum* 30% alcohol extract, *Cryptolepis sanguinolenta* 30%, 60%, and 90% alcohol extracts, *Scutellaria baicalensis* (huang qin), and *Scutellaria barbata* (ban zhi lian) had similar activity against growing *B*. *henselae* such that they inhibited log phase *B. henselae* proliferation at 0.25%-0.5% (*v/v*). These results indicated that these top hits of herbal products were not only active against non-growing stationary phase *B. henselae*, but also effective against log phase growing *B. henselae.*

**Table 3.**
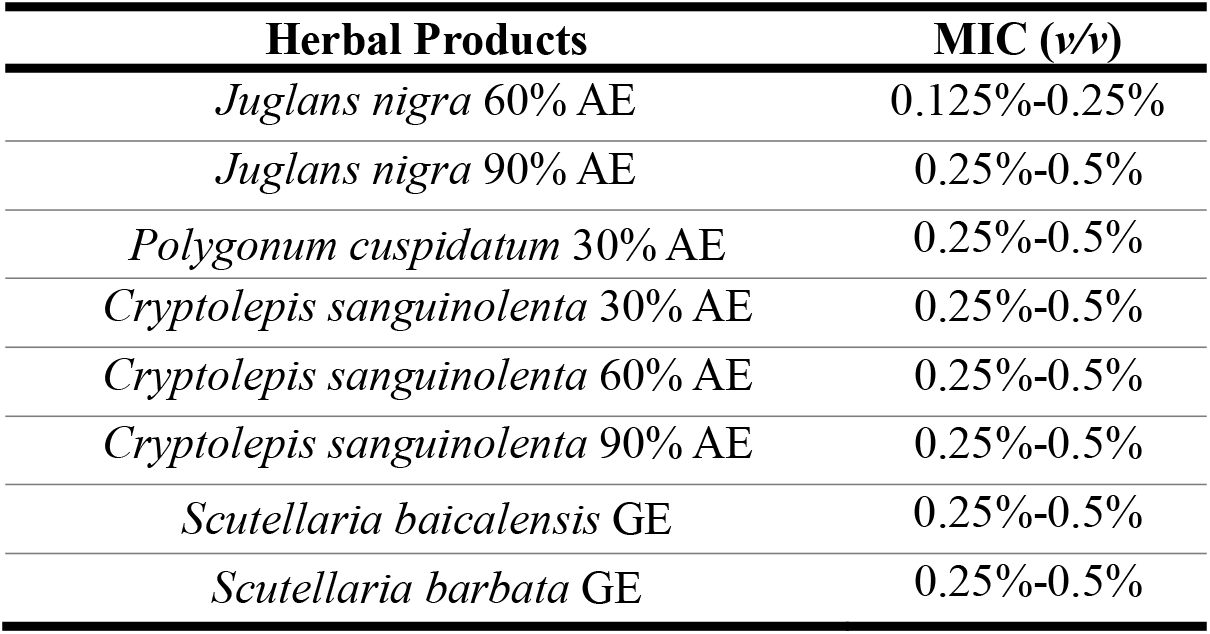
Minimum inhibitory concentrations (MICs) of top active herbal products against *B. henselae*.

| Herbal Products | MIC (v/v) |
| --- | --- |
| *Juglans nigra* 60% AE | 0.125%-0.25% |
| *Juglans nigra* 90% AE | 0.25%-0.5% |
| *Polygonum cuspidatum* 30% AE | 0.25%-0.5% |
| *Cryptolepis sanguinolenta* 30% AE | 0.25%-0.5% |
| *Cryptolepis sanguinolenta* 60% AE | 0.25%-0.5% |
| *Cryptolepis sanguinolenta* 90% AE | 0.25%-0.5% |
| *Scutellaria baicalensis* GE | 0.25%-0.5% |
| *Scutellaria barbata* GE | 0.25%-0.5% |

## DISCUSSION

In this study, we successfully applied the SYBR Green I/PI viability assay to evaluate a panel of botanical products for their activity against stationary phase *B. henselae* as a model of persister drug screens [51][57]. We identified some herbal products that had high activity at 1% (*v/v*) concentration compared to clinically used antibiotics, including extracts of *Juglans nigra*, *Cryptolepis sanguinolenta*, *Polygonum cuspidatum*, *Scutellaria baicalensis*, and *Scutellaria barbata*. Among these top hits, three herbal product extracts could eradicate all stationary *B. henselae* cells without CFU being detected within a seven-day drug exposure at a low concentration of 0.25% (*v/v*), including *Cryptolepis sanguinolenta* 30%, 60%, 90% alcohol extracts, *Juglans nigra* 60%, 90% alcohol extracts, and *Polygonum cuspidatum* 30% alcohol extracts. The MIC determination of these active hits showed they were also effective in inhibiting the growth of log phase *B. henselae*.

These plant species whose extracts we found to be active against *B. henselae* have also been reported to have various biological activities in previous studies. Different parts of various species from genus *Juglans* have shown pain-relieving, antioxidant, antibacterial, antifungal and antitumor activities [66–68]. In particular, *Juglans nigra* exhibited both bacteriostatic activity and bactericidal activity against *Borrelia* based on *in vitro* studies [60][69]. Previous studies have profiled the phytochemicals of *Juglans* plants, including different types of steroids, flavonoid C-glycoside, flavones, essential oil components, and tannins [70]. *Juglans* contain several active constituents with potential importance to human health including juglone, phenolic acids, flavonoids, and catechins (including epigallocatechin) [71–75]. A study comparing leaf essential oils of *J. regia* and *J. nigra* further showed *J. nigra* leaf oil was less phytotoxic [76]. The safety of *Juglans nigra* use in humans has not been adequately studied however it has a long history of anecdotal use and the oral LD50 of juglone in rats is calculated at 112mg/kg [77].

*Cryptolepis sanguinolenta* and its constituents were reported to have many biological activities including antibacterial, antifungal, anti-inflammatory, anticancer, antimalarial and anti-amoebic properties [61][78–82]. A recent review has assessed the phytochemistry and pharmacology of *Cryptolepis sanguinolenta* and concluded that although there may be some concern regarding potential reproductive toxicity, it is generally safe at doses below 500mg/kg and may serve as promising source of potential antimicrobial agent(s) [83]. Among constituents and secondary metabolites of the plant identified with antimicrobial activity, an alkaloid called cryptolepine was the most well-studied and considered to be the most important active component. Cryptolepine was reported to have a lytic effect on *S. aureus* as seen in SEM photomicrographs which led to altered cell morphology, and was able to intercalate into DNA at cytosine-cytosine sites or inhibited the activity of topoisomerase causing DNA damage [84–86]. *Cryptolepis sanguinolenta* was reported to be highly effective and non-toxic in the treatment of uncomplicated malaria in a small randomized open trial of 44 patients [87]. *Cryptolepis sanguinolenta* is also used anecdotally to treat a malaria-like tickborne infection called Babesiosis [88]. Given its anecdotal use to treat Babesiosis, the results of the current study on *B. henselae,* and the results of our previous study on *B. burgdorferi* [56]*, Cryptolepis sanguinolenta* represents a unique potential therapeutic agent to treat multiple tickborne infections. Future studies are needed to elucidate more specific antimicrobial mechanisms of cryptolepine as well as other active ingredients against infectious pathogens such as *B. henselae*. *Polygonum cuspidatum* has been documented to have antibacterial effects against *Vibrio vulnificus* [89], *Streptococcus mutans* [90] and *Streptococcus* associated biofilms [91]. Its constituents have also been shown to have antimicrobial, anti-tumor, anti-inflammatory, neuroprotective, and cardioprotective effects [92–96]. One of the most active constituents is a polyphenol called resveratrol, which was reported to be active against log phase *Borrelia burgdorferi* and *Borrelia garinii* by *in vitro* testing [60]. In addition, another active constituent called emodin (6-methyl-1,3,8-trihydroxyanthraquinone) has been shown to have activity against stationary phase *Borrelia burgdorferi* cells [97]. A study unraveling the mechanism of action of *Polygonum cuspidatum* using a network pharmacology approach, suggested that polydatin might play a pivotal role in the therapeutic effects of *Polygonum cuspidatum* [98]. Recent trials using *Polygonum cuspidatum* have not reported significant toxicity [99, 100]. One of the active constituents, resveratrol, has been shown to be rapidly absorbed [101], well-tolerated [102] and associated with minimal toxicity except mild diarrhea at an oral dose of 2000mg 2x/day for 2 weeks [103].

Our study is the first to identify the antimicrobial activity of extracts from *Juglans nigra*, *Cryptolepis sanguinolenta*, and *Polygonum cuspidatum* against stationary phase *B. henselae.* In addition, considering the possibility of *B. henselae* coinfection among Lyme and tick-borne disease patients, the overlap of active herbal products against both *B. henselae* identified in our current study and *B. burgdorferi* according to our previous study [56], including *Cryptolepis sanguinolenta, Juglans nigra*, and *Polygonum cuspidatum*, should provide a promising strategy for better treatment of coinfections with both pathogens.

In this current study, clinically used antibiotics for treating *Bartonella*-associated infections including AZI and DOX showed weak activity in eradicating stationary phase *B. henselae* cells (Table 1, Figure 1 and Figure 2). This finding coincides with the reported discrepancies in antibiotic efficacies between *in vitro* MIC data and clinical data from patients [35]. The poor activities of current clinically used antibiotics against stationary phase *B. henselae* as shown in our study could partly explain clinically documented treatment failure and could be in part due to persistent infection. This phenomenon may also be partly due to the limited antibacterial activity of these antibiotics. Doxycycline inhibits bacterial protein synthesis by binding to the 30S ribosomal subunit [104]. Azithromycin could also inhibit bacterial protein synthesis by binding to the 50S ribosomal subunit, and thus prevent bacteria from growing [105]. Although these antibiotics all target growing bacteria, they are not very effective at killing non-growing stationary phase *B. henselae*, and thus could lead to treatment failure in persistent and chronic infections. Conversely, the herbal extracts could be promising candidates for treating persistent *B. henselae* because they contain multiple active phytochemicals including steroids, flavones, tannins and more. These compounds have complex and synergistic effects and thus have potentially broader antimicrobial activity. Many of these phytochemicals are lipophilic and could target the bacterial cell membrane, which is an important target of persister drugs like pyrazinamide [106] and daptomycin [107], especially when persistent bacterial cells are aggregated together (Figure 1). The high lipophilicity of these phytochemicals, which could cause bacterial cell membrane damage, could be responsible for the aggregated bacterial forms and also explain for the varying numbers of bacterial cells among different samples seen in microscopic pictures (Figure 1).

Despite the promising findings of the herbal extracts active against *B. henselae*, future studies are needed to identify the active ingredients of these herbs and to better understand their specific antimicrobial mechanisms of action. It would also be of interest to test compounds like juglone, cryptolepine, resveratrol, and emodin which are known active components of the *Juglans nigra*, *Cryptolepis sanguinolenta* and *Polygonum cuspidatum* herbs, on *B. henselae* in future studies. Different parts of these plants might have different antimicrobial activities because of varying concentrations of the active compounds they contained, and different solvents used to extract the compounds could also significantly affect their activity. Therefore, the pharmacokinetic profiles of active components should be studied thoroughly in the future, as well as the optimal extraction strategy to obtain the maximally effective ingredients in order to better determine the utility and practicality of these active herbal medicines.

The present study only tested the activity of herbal products against *B. henselae in vitro* and there are a few points that are important to address. For one, *B. henselae* is a facultative intracellular pathogen and could reside and propagate inside mammalian erythrocytes and/or endothelial cells. Therefore, it will be of interest to assess the activity of these identified candidates against intracellular *B. henselae ex vivo* and *in vivo* in animal models of infection [108] in the future. The host cell can provide the pathogen with protective shelter from the action of drugs and herbal antimicrobials as well as the host immune system, therefore the efficacy of these antimicrobials *in vivo* might differ from that *in vitro* [11]. Additionally, it is important to note that botanical medicines have multiple mechanisms of action beyond the antimicrobial activity that was assessed in the present study. For example, botanicals have been shown to exert effects via multiple mechanisms with potential benefit in *Bartonella* infections including anti-inflammatory activity, immune modulation/stimulation, microbiome modulation, endothelial cell support and biofilm disruption. Future studies are needed to assess the safety and efficacy of these herbal products in appropriate animal models of *Bartonella* infections where broader biologic mechanisms of action including their effects on the host can be evaluated.

In the future we also hope to test different combinations of active herbal products and their active constituents with and without antibiotics to develop better treatments. As indicated by the Yin-Yang model, the bacterial pathogen has a heterogeneous population, with persister population (Yin) and growing population (Yang), which are also composed of various subpopulations with varying metabolic or dormant states in continuum [38]. Therefore, it should be reasonable to deploy drug/herb or herb/herb combinations for more effective treatments, with different drugs or herbal medicines targeting different bacterial subpopulations in varying physiological states. Indeed, members of our group recently demonstrated that antibiotic combinations were more effective in eradicating *in vitro* stationary phase and biofilm *B. henselae* compared to single antibiotics [109]. Our goal is to use the herbal medicines we identified in this study to develop more safe and effective treatments for persistent bartonellosis.

## Acknowledgments

We thank herbalists Eric Yarnell ND and Brian Kie Weissbuch for providing botanical extracts for evaluation in this study. We thank BEI Resources/ATCC, NIAID, NIH for providing *Bartonella henselae* JK53 strain used in this study. We gratefully acknowledge the support of our work by the Bay Area Lyme Foundation, the Steven & Alexandra Cohen Foundation, LivLyme Foundation, Global Lyme Alliance, NatCapLyme, and the Einstein-Sim Family Charitable Fund.

---

**Table S1.**
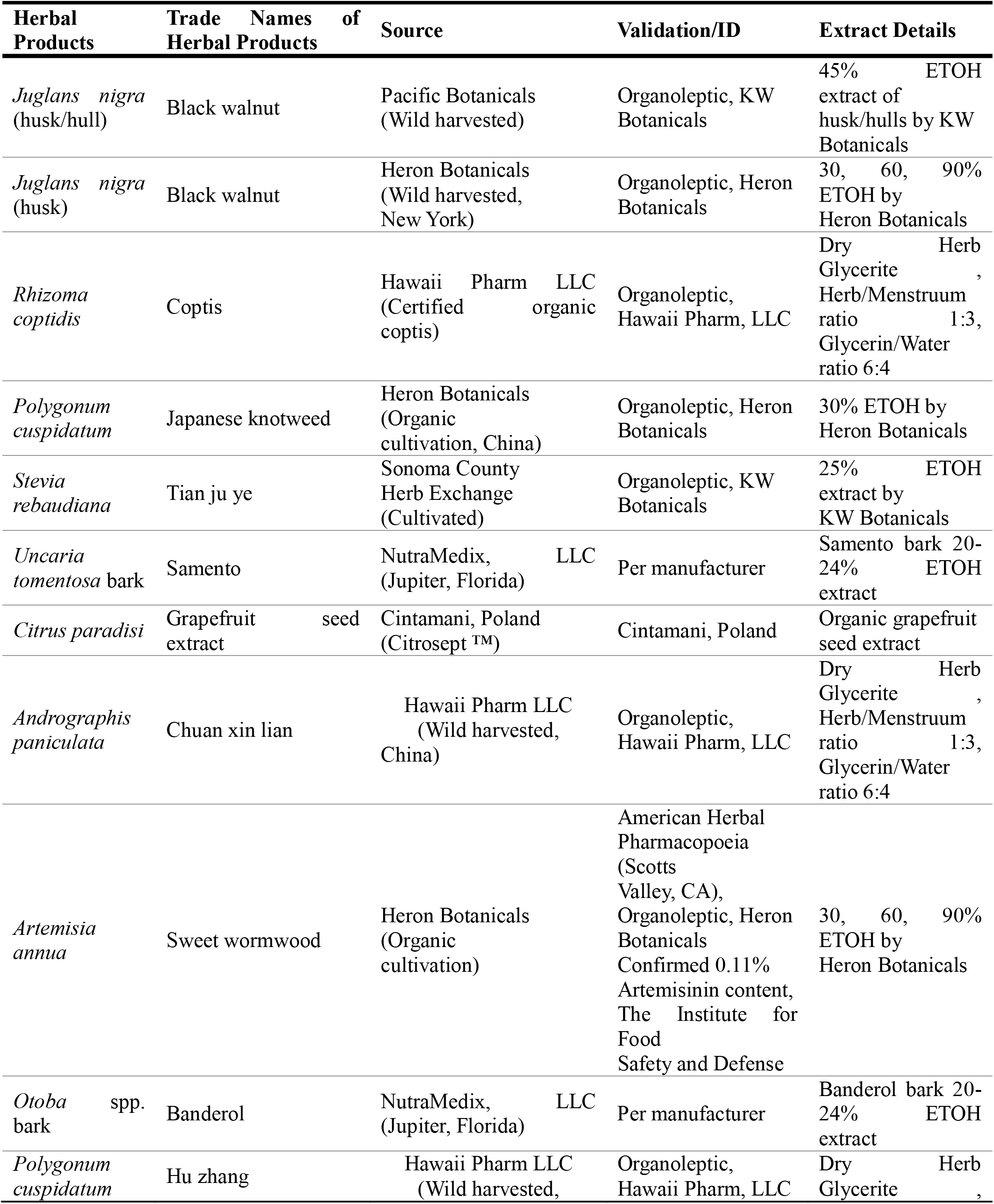
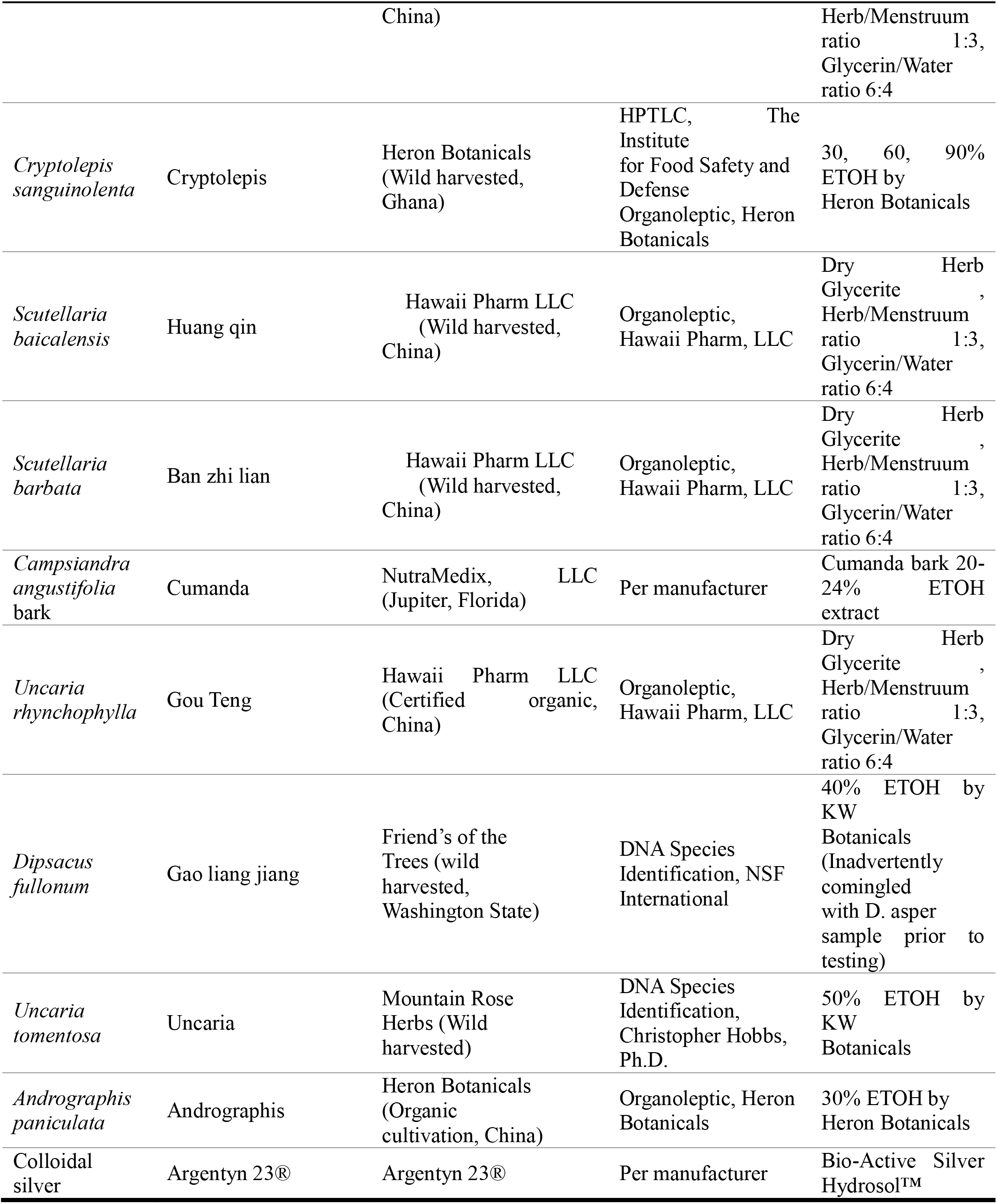
Herbal product sources, validation, and extract details.

**Table S2.**
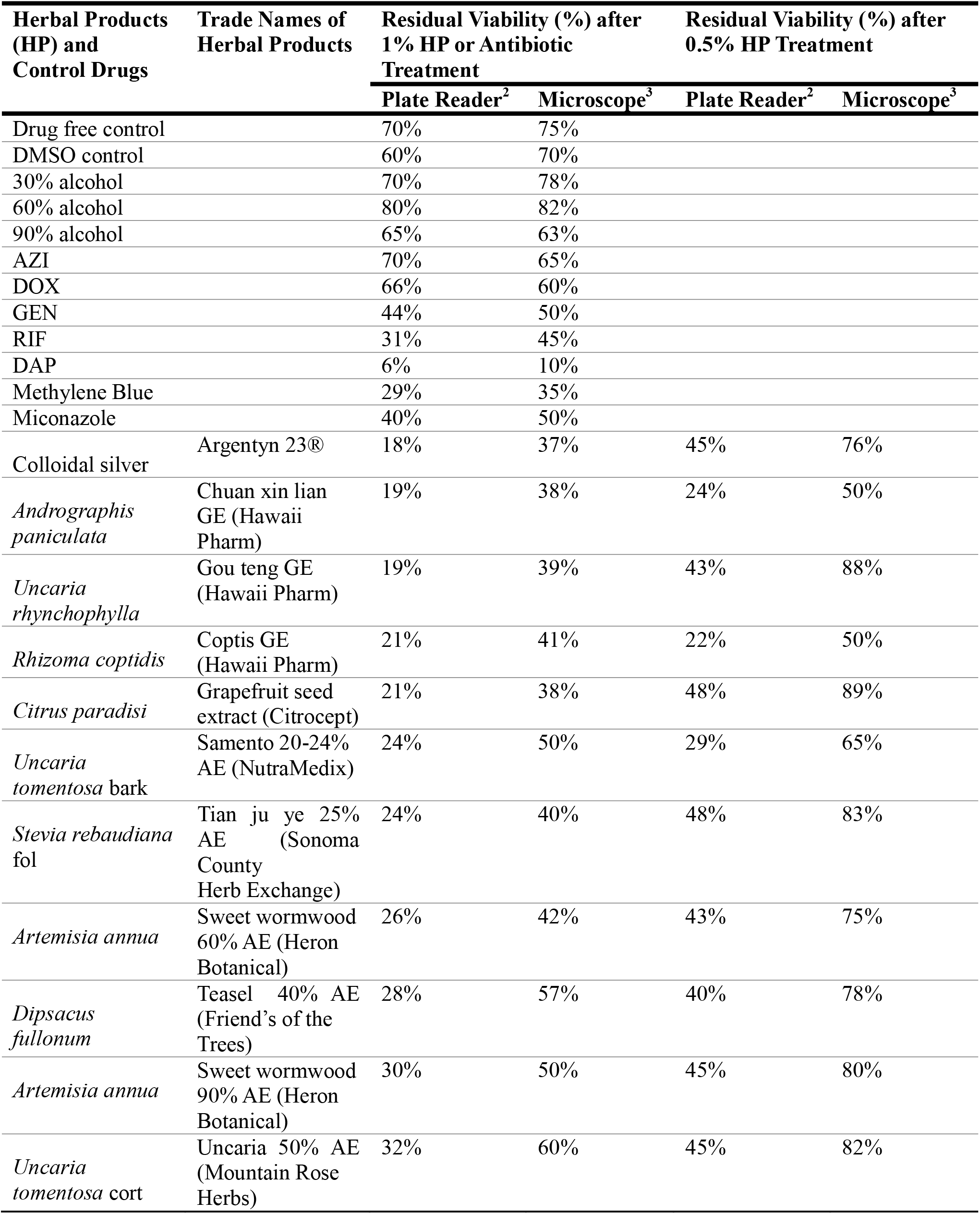
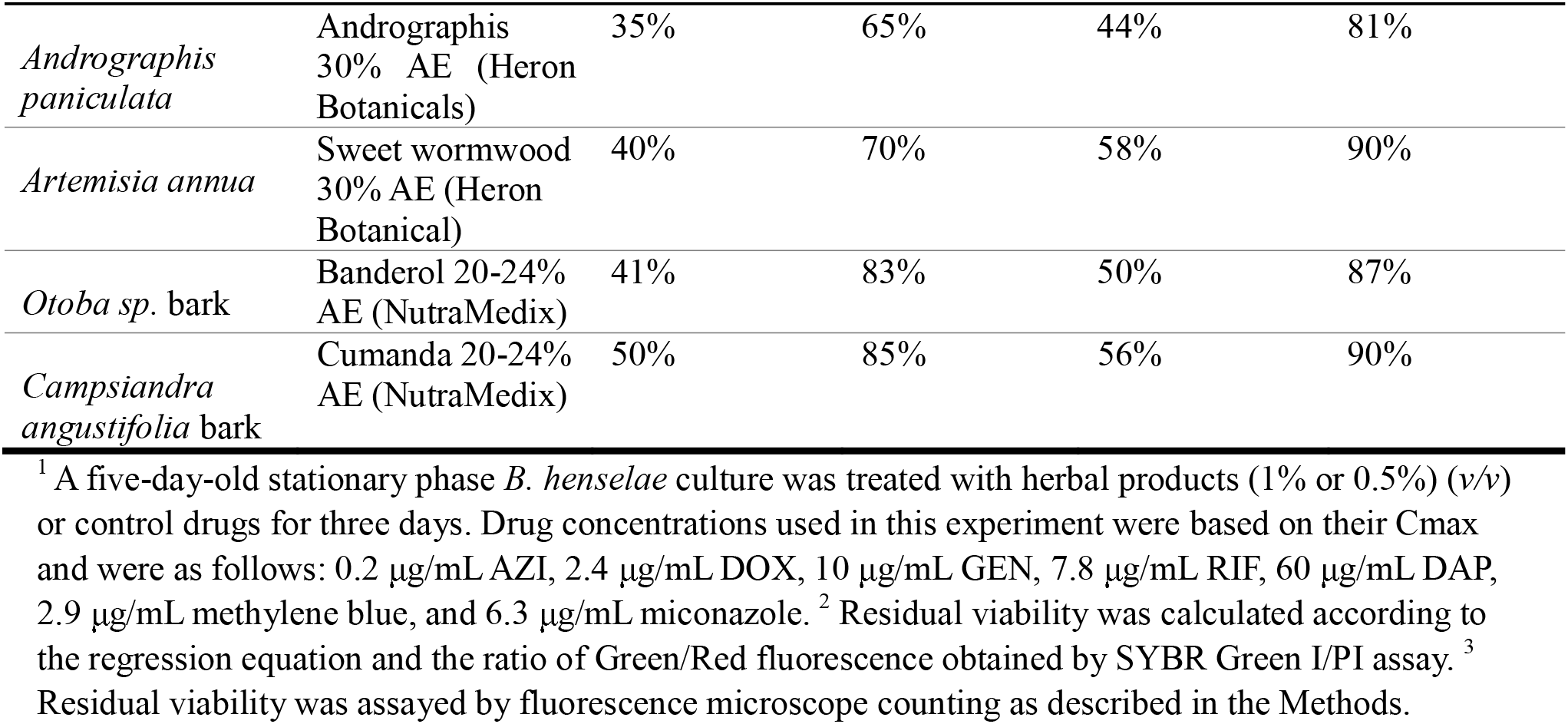
Activity of other tested herbal products against stationary phase *B. henselae* ^1^.

