## Supplementary material for "Botanical Medicines with Activity against Stationary Phase *Bartonella henselae*": Table S1 S2

**Table S1. Herbal product sources, validation, and extract details**

| **Herbal Products** | **Trade Names of Herbal Products** | **Source** | **Validation/ID** | **Extract Details** |
| --- | --- | --- | --- | --- |
| *Juglans nigra* (husk/hull) | Black walnut | Pacific Botanicals  (Wild harvested) | Organoleptic, KW  Botanicals | 45% ETOH extract of  husk/hulls by KW  Botanicals |
| *Juglans nigra* (husk) | Black walnut | Heron Botanicals  (Wild harvested,  New York) | Organoleptic, Heron  Botanicals | 30, 60, 90% ETOH by  Heron Botanicals |
| *Rhizoma coptidis* | Coptis | Hawaii Pharm LLC (Certified organic coptis) | Organoleptic, Hawaii Pharm, LLC | Dry Herb Glycerite , Herb/Menstruum ratio 1:3, Glycerin/Water ratio 6:4 |
| *Polygonum cuspidatum* | Japanese knotweed | Heron Botanicals  (Organic  cultivation, China) | Organoleptic, Heron  Botanicals | 30% ETOH by  Heron Botanicals |
| *Stevia rebaudiana* | Tian ju ye | Sonoma County  Herb Exchange  (Cultivated) | Organoleptic, KW  Botanicals | 25% ETOH extract by  KW Botanicals |
| *Uncaria tomentosa* bark | Samento | NutraMedix, LLC (Jupiter, Florida) | Per manufacturer | Samento bark 20-24% ETOH extract |
| *Citrus paradisi* | Grapefruit seed extract | Cintamani, Poland  (Citrosept ™) | Cintamani, Poland | Organic grapefruit  seed extract |
| *Andrographis paniculata* | Chuan xin lian | Hawaii Pharm LLC (Wild harvested,  China) | Organoleptic, Hawaii Pharm, LLC | Dry Herb Glycerite , Herb/Menstruum ratio 1:3, Glycerin/Water ratio 6:4 |
| *Artemisia annua* | Sweet wormwood | Heron Botanicals  (Organic  cultivation) | American Herbal  Pharmacopoeia (Scotts  Valley, CA),  Organoleptic, Heron  Botanicals  Confirmed 0.11%  Artemisinin content,  The Institute for Food  Safety and Defense | 30, 60, 90% ETOH by  Heron Botanicals |
| *Otoba* spp. bark | Banderol | NutraMedix, LLC (Jupiter, Florida) | Per manufacturer | Banderol bark 20-24% ETOH extract |
| *Polygonum cuspidatum* | Hu zhang | Hawaii Pharm LLC (Wild harvested,  China) | Organoleptic, Hawaii Pharm, LLC | Dry Herb Glycerite , Herb/Menstruum ratio 1:3, Glycerin/Water ratio 6:4 |
| *Cryptolepis sanguinolenta* | Cryptolepis | Heron Botanicals  (Wild harvested,  Ghana) | HPTLC, The Institute  for Food Safety and  Defense  Organoleptic, Heron  Botanicals | 30, 60, 90% ETOH by  Heron Botanicals |
| *Scutellaria baicalensis* | Huang qin | Hawaii Pharm LLC (Wild harvested,  China) | Organoleptic, Hawaii Pharm, LLC | Dry Herb Glycerite , Herb/Menstruum ratio 1:3, Glycerin/Water ratio 6:4 |
| *Scutellaria barbata* | Ban zhi lian | Hawaii Pharm LLC (Wild harvested,  China) | Organoleptic, Hawaii Pharm, LLC | Dry Herb Glycerite , Herb/Menstruum ratio 1:3, Glycerin/Water ratio 6:4 |
| *Campsiandra angustifolia* bark | Cumanda | NutraMedix, LLC (Jupiter, Florida) | Per manufacturer | Cumanda bark 20-24% ETOH extract |
| *Uncaria rhynchophylla* | Gou Teng | Hawaii Pharm LLC (Certified organic, China) | Organoleptic, Hawaii Pharm, LLC | Dry Herb Glycerite , Herb/Menstruum ratio 1:3, Glycerin/Water ratio 6:4 |
| *Dipsacus fullonum* | Gao liang jiang | Friend’s of the  Trees (wild  harvested,  Washington State) | DNA Species  Identification, NSF  International | 40% ETOH by KW  Botanicals  (Inadvertently comingled  with D. asper  sample prior to testing) |
| *Uncaria tomentosa* | Uncaria | Mountain Rose  Herbs (Wild  harvested) | DNA Species  Identification,  Christopher Hobbs,  Ph.D. | 50% ETOH by KW  Botanicals |
| *Andrographis paniculata* | Andrographis | Heron Botanicals  (Organic  cultivation, China) | Organoleptic, Heron  Botanicals | 30% ETOH by  Heron Botanicals |
| Colloidal silver | Argentyn 23® | Argentyn 23® | Per manufacturer | Bio-Active Silver Hydrosol™ |

**Table S2. Activity of other tested herbal products against stationary phase *B. henselae* ^1^**

| **Herbal Products (HP) and Control Drugs** | **Trade Names of Herbal Products** | **Residual Viability (%) after 1% HP or Antibiotic Treatment** | | **Residual Viability (%) after 0.5% HP Treatment** | |
| --- | --- | --- | --- | --- | --- |
|  |  | **Plate Reader^2^** | **Microscope^3^** | **Plate Reader^2^** | **Microscope^3^** |
| Drug free control |  | 70% | 75% |  |  |
| DMSO control |  | 60% | 70% |  |  |
| 30% alcohol |  | 70% | 78% |  |  |
| 60% alcohol |  | 80% | 82% |  |  |
| 90% alcohol |  | 65% | 63% |  |  |
| AZI |  | 70% | 65% |  |  |
| DOX |  | 66% | 60% |  |  |
| GEN |  | 44% | 50% |  |  |
| RIF |  | 31% | 45% |  |  |
| DAP |  | 6% | 10% |  |  |
| Methylene Blue |  | 29% | 35% |  |  |
| Miconazole |  | 40% | 50% |  |  |
| Colloidal silver | Argentyn 23® | 18% | 37% | 45% | 76% |
| *Andrographis paniculata* | Chuan xin lian GE (Hawaii Pharm) | 19% | 38% | 24% | 50% |
| *Uncaria rhynchophylla* | Gou teng GE (Hawaii Pharm) | 19% | 39% | 43% | 88% |
| *Rhizoma coptidis* | Coptis GE (Hawaii Pharm) | 21% | 41% | 22% | 50% |
| *Citrus paradisi* | Grapefruit seed extract (Citrocept) | 21% | 38% | 48% | 89% |
| *Uncaria tomentosa* bark | Samento 20-24% AE (NutraMedix) | 24% | 50% | 29% | 65% |
| *Stevia rebaudiana* fol | Tian ju ye 25% AE (Sonoma County  Herb Exchange) | 24% | 40% | 48% | 83% |
| *Artemisia annua* | Sweet wormwood 60% AE (Heron Botanical) | 26% | 42% | 43% | 75% |
| *Dipsacus fullonum* | Teasel 40% AE (Friend’s of the  Trees) | 28% | 57% | 40% | 78% |
| *Artemisia annua* | Sweet wormwood 90% AE (Heron Botanical) | 30% | 50% | 45% | 80% |
| *Uncaria tomentosa* cort | Uncaria 50% AE (Mountain Rose  Herbs) | 32% | 60% | 45% | 82% |
| *Andrographis paniculata* | Andrographis 30% AE (Heron Botanicals) | 35% | 65% | 44% | 81% |
| *Artemisia annua* | Sweet wormwood 30% AE (Heron Botanical) | 40% | 70% | 58% | 90% |
| *Otoba sp.* bark | Banderol 20-24% AE (NutraMedix) | 41% | 83% | 50% | 87% |
| *Campsiandra angustifolia* bark | Cumanda 20-24% AE (NutraMedix) | 50% | 85% | 56% | 90% |

^1^ A five-day-old stationary phase *B. henselae* culture was treated with herbal products (1% or 0.5%) (*v/v*) or control drugs for three days. Drug concentrations used in this experiment were based on their Cmax and were as follows: 0.2 μg/mL AZI, 2.4 μg/mL DOX, 10 μg/mL GEN, 7.8 μg/mL RIF, 60 μg/mL DAP, 2.9 μg/mL methylene blue, and 6.3 μg/mL miconazole. ^2^ Residual viability was calculated according to the regression equation and the ratio of Green/Red fluorescence obtained by SYBR Green I/PI assay. ^3^ Residual viability was assayed by fluorescence microscope counting as described in the Methods.
